# Evidence for intercellular bridges and radial patterning of meiotic initiation in the human fetal ovary

**DOI:** 10.1101/2022.07.18.500281

**Authors:** Bikem Soygur, Amber Derpinghaus, Gerald R. Cunha, Laurence S. Baskin, Diana J Laird

## Abstract

Meiosis is the hallmark of reproduction. Our understanding of early oocyte development was improved by studying the spatiotemporal dynamics and mechanisms governing meiosis in mice, however, our knowledge of the meiotic initiation process in humans remains limited. Here, we utilized three-dimensional (3D) analysis to determine spatiotemporal dynamics of meiotic initiation in fetal human ovaries. Similar to mice, we found that the first meiotic cells appear in clusters in the center of human fetal ovaries as early at 9 weeks and that the initiation of meiosis propagates as a radial wave. Between developing germ cells in fetal human ovaries, we detected a component of the intercellular bridge, TEX14 protein. 3D quantification of germ cells in ovaries collected at the end of first trimester revealed, for the first time, considerable variation in the number of meiotic cells between individuals and asynchronous mitotic-meiotic transition. In addition to illustrating the geography of meiotic initiation in ovaries from the first trimester, we extended our 3D analysis approach to second trimester and showed heterogeneous spatial distribution of meiotic and non-meiotic germ cells in human fetal ovaries. This study is an important step towards better understanding of 3D structure of developing ovary and early stages of meiosis in humans.

**Highlights:**

- Organic solvent-based clearing methods and confocal microscopy can be implemented to visualize and quantify germ cells in the intact human fetal ovary.
- Identification of a new, radial, pattern of meiotic initiation in the ovaries from the first trimester.
- Immunolocalization of the intercellular bridge component TEX14 between developing germ cells in the fetal ovary
- Maintenance of pre-meiotic germ cells in second trimester ovaries suggests less synchronous mitotic-meiotic stage transition.

## Introduction

Gamete precursors were first detected in 3-week old human embryos near the yolk sac wall in 1911 (Felix, 1911; Fuss, 1911). More than century later, although much is known about primordial germ cell (PGC) development, the precise timing and location of germline specification in humans remains unclear owing to the technical and ethical difficulties of studying human embryos [reviewed in (Hancock, Wamaitha, Peretz, & Clark, 2021; Kobayashi & Surani, 2018)]. Recently, bioinformatic analysis of a gastrulation stage (CS7) human embryo revealed a small population of primitive streak cells expressing high levels of PGC markers, suggesting the presence of PGCs as early as 16-19 days after fertilization (Tyser et al., 2021). Following their specification, PGCs migrate and reach the gonads starting at the sixth week (Makabe, Naguro, Nottola, Pereda, & Motta, 1991; Mc, Hertig, Adams, & Danziger, 1953; Witschi, 1948) where they proliferate massively. During the proliferative stage, stable intercellular bridges are formed between germ cells; however, their function in developing human ovary is unknown [reviewed in (Greenbaum, Iwamori, Buchold, & Matzuk, 2011)]. Gamete differentiation of the XX embryo, begins during the first trimester soon after PGCs settle down in the gonads with the initiation of meiosis, which occurs asynchronously over many weeks (Baker, 1963; Kurilo, 1981; Skrzypczak, Pisarski, Biczysko, & Kedzia, 1981); this transition from mitotic germ cells (oogonia) to meiotic oocytes is one of the earliest and the major commitments of female germ cell differentiation. As mitotic divisions cease after meiosis begins in fetal human ovaries, the number of germ cells and their developmental trajectory during fetal development are an indicator of future reproductive potential.

Although recently developed imaging and tissue clearing methods have advanced the study of meiotic initiation in mice (Soygur et al., 2021), our understanding of early development of female germ cells in humans is largely based on cytological (Baker, 1963; Manotaya & Potter, 1963), histological (Bendsen, Byskov, Andersen, & Westergaard, 2006; Kurilo, 1981), and electron microscopy techniques (Gondos, Bhiraleus, & Hobel, 1971; Gondos, Westergaard, & Byskov, 1986). Chromosomal and light microscopic analyses revealed the presence of preleptotene stage germ cells that are characterized by dense chromosome masses in human ovaries from as early as 9-10 weeks of fetal development (fetal age; FA) (Bendsen et al., 2006; Gondos et al., 1986). Molecular and immunohistochemical studies later reported that transcripts (*STRA8, SPO11*, and *DMC1*) and protein (γH_2_AX) associated with meiosis were first detected in 11-week-old fetal ovaries (Le Bouffant et al., 2010). While meiotic initiation in human fetal ovaries has been described in numerous studies, the results vary, and our understanding of early germ cell development is incomplete. Discrepancies between studies arise from differences in embryo staging or from the challenges of using the sampling method, which is labor intensive and can easily miss rare developmental events.

Recently emerging whole-mount imaging and analysis technologies provide three-dimensional (3D) information at the cellular level using intact tissues and organs. Such holistic analysis at single cell resolution holds the capacity to decode embryogenesis, organogenesis, and ultimately disease. Studies in mice highlighted the importance of 3D analysis of developing embryos, organs, and even a whole adult mouse body (Pan et al., 2016) by unveiling new cell populations or spatiotemporal regulations that have not been described with earlier sampling methods [reviewed in (Molbay, Kolabas, Todorov, Ohn, & Erturk, 2021)]. However, translating the techniques and methods pioneered in mouse to human has proved non-trivial, due to the increased complexity of human tissue, both in size and architecture, as well as the limited availability of human embryonic/fetal samples. To date, the most comprehensive 3D map of the human development includes visualization of embryos/fetuses from 6 to 14-weeks (Belle et al., 2017). Although successful 3D rendering was performed on various organs and organ systems including the genital ducts of urogenital system, germ cell development in developing ovaries has not been investigated.

Here we employed whole-mount immunofluorescence staining in combination with confocal imaging and 3D analysis to visualize and quantify different germ cell populations in developing human fetal ovaries. Employing several markers to distinguish different stages of germ cell development, we investigated the spatiotemporal dynamics of the mitotic-meiotic transition in human fetal ovaries *in toto*. This powerful 3D analysis method revealed that germ cells initiate meiosis as early as 9-weeks with the first meiotic cells appearing in the center of the ovary. This radial pattern of meiotic entry was observed in all samples analyzed in the first trimester. We showed that meiosis initiates in germ cell clusters (nests) in human fetal ovaries, as occurs in mice but is less synchronous in humans; coincident detection of the intercellular bridge component TEX14 raises the possibility that the timing of meiotic entry involves intercellular bridges between germ cells in humans as in mice (Soygur et al., 2021).

## Materials and Methods

### Sample Collection

Following elective termination procedures, human fetal ovaries were collected with the approval of the institutional review board at the University of California, San Francisco (UCSF 16–19909 #167670). The collection technique was previously described (Shen et al., 2018). Heel-toe length was used to determine the age of specimens (Drey, Kang, McFarland, & Darney, 2005). Specimen sex was determined by morphology of the testes or ovaries and/or morphology of the Wolffian and Mullerian ducts. Dissected fetal ovaries were kept in Hank’s Balanced Salt Solution (HBSS with calcium and magnesium; Thermo Fisher Scientific, cat# 24020117) at 4°C for several hours preceding fixation.

### Whole-mount immunofluorescence

Fetal ovaries were transferred to 2-ml Eppendorf tubes and fixed with 4% paraformaldehyde (PFA) in PBS on an orbital shaker at 4°C for 4 hours (h). All subsequent incubation steps were carried with rocking. Ovaries were washed three times with PBS for 20 minutes (mins) each. We used two different organic solvent-based clearing agents to achieve transparency as previously described (Soygur et al., 2022). Fetal ovaries were blocked with blocking buffer for 4 hours at room temperature. Primary antibodies (OCT4; Santa Cruz #sc-5279, VASA; Novus Biologicals #AF2030, SYCP3; Novus Biologicals #NB300-232, ɣH2AX; Millipore #05636) were diluted in antibody dilution solution, and ovaries were incubated in primary antibodies at 4°C for a week. Samples were washed three times with washing solution for 20 min each at room temperature and incubated with Alexa Fluor–conjugated secondary antibodies in antibody dilution solution at 4°C for 5 days. Ovaries were washed three times with washing solution for 20 min each, dehydrated with methanol:PBS series (25 to 50 to 75 to 100%) for 20 min each (only 100% twice) at room temperature The following day, ovaries were incubated in organic solvent-based clearing solution, transferred to the 10-mm-long glass cylinders (ACE Glass 3865-10) mounted onto coverslips (Fisherfinest Premium Cover Glass 12–548-5P).

Samples were imaged with a white-light Leica TCS SP8 converted confocal microscope with a HC PL APO CS 10X∼/0.40 dry objective or a Fluotar VISIR 25X/0.95 water objective, 0.75X optic zoom, and 1024 X1024 pixel resolution.

### Section Immunofluorescence

Samples were fixed with 2% PFA in PBS for overnight, washed four times with PBS for 20 mins each and dehydrated overnight in 30% sucrose at 4°C. The ovaries were embedded in optimal cutting temperature compound, flash-frozen and stored at −80°C. Cryostat sections were cut at a thickness of 12 μm and affixed to Superfrost Plus slides (Fisher Scientific). The sections were washed three times with PBS for 10 mins each, blocked with 5% donkey serum, 1% Triton X-100 in PBS for 1 h at room temperature and incubated overnight at 4°C with the primary antibodies (VASA; AF2030, TEX14; ab41733) in blocking solution. The next day, the sections were washed with PBS for 10 mins three times and incubated with Alexa Fluor–conjugated secondary antibodies for 1 h at room temperature. The slides were washed with PBS for 10 mins three times, mounted with VECTASHIELD, and imaged on a SP5 Leica confocal microscope.

### Image Analysis for whole-mount stained ovaries

Image analysis was performed using Imaris v9.7.1 (Bitplane). The files were converted with Imaris converter v9.7 and imported to Surpass mode. The background subtraction value=8 was applied to the samples imaged with a 10X objective to increase the accuracy of the quantification. Fluorescently labeled germ cells were selected by using the Spot detection module; XY diameter of 4 μm (object size) were applied to detect labeled germ cells. For the samples imaged with a 25X objective, the background subtraction value was determined as 4.75 and XY diameter of 4 μm spots were selected to detect germ cells. Germ cell numbers for each marker were imported to a spreadsheet application.

## Results

### Developing methods for 3D analysis of meiotic initiation in human fetal ovaries

To visualize germ cells in intact ovaries, we performed whole-mount immunofluorescent staining using antibodies against markers of non-meiotic and meiotic germ cells in fetal human ovaries (**Figure 1A**). Following organic solvent-based clearing, fetal ovaries were imaged by confocal fluorescent microscopy. Consistent with previous 2D histological analysis (Anderson, Fulton, Cowan, Coutts, & Saunders, 2007), the majority of germ cells expressed OCT4 (marker for pluripotent germ cells) in ovaries at 9 weeks of fetal age (FA; **Figure 1B**); with our 3D analysis strategy, we quantified more than 100,000 OCT4 positive cells (OCT4+) per ovary. While a subset of germ cells expressed VASA (marker for developmentally-progressed germ cells), co-expression of SYCP3 (marker for meiotic germ cells) was detected in vanishingly few germ cells (12 cells; only 0.0089% of the total population), indicating that meiosis begins as early as 9 weeks in fetal human ovaries (**Figure 1B; higher magnification and Figure 2C**). 3D reconstruction of the entire ovary revealed that the earliest SYCP3 expressing cells were located in the central region of the human fetal ovary, near the mesonephric-gonadal junction, and they tended to group together (**Figure 1C**).

**Figure 1.**
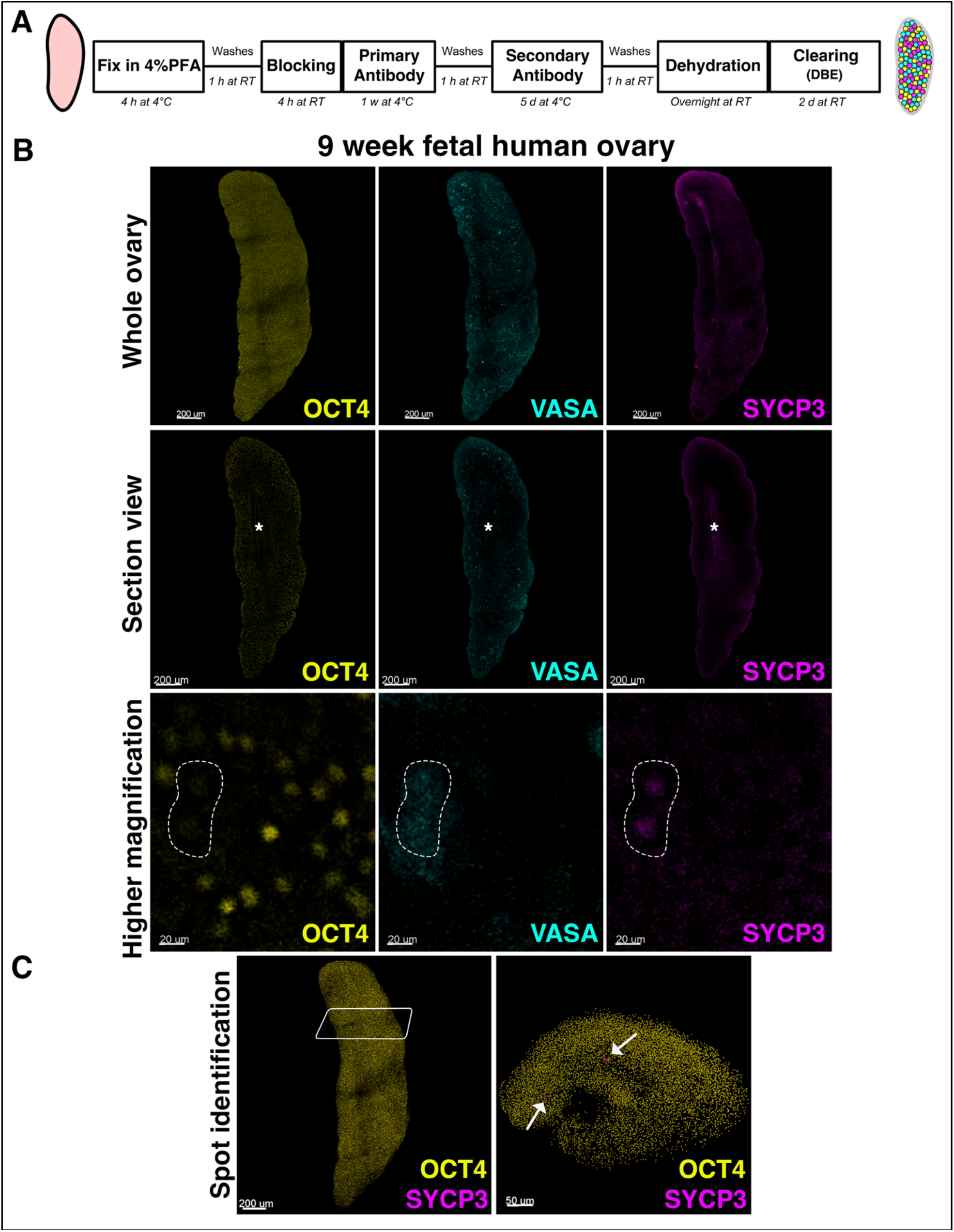
3D whole-mount analysis of the human fetal ovary. **A**. Flowchart of whole-mount immunofluorescence labeling and organic solvent-based clearing (using DBE) procedures of human fetal ovaries. Dark lines are artifacts of image stitching. **B**. An ovary from 9 weeks FA stained with OCT4 (pluripotency marker in yellow), VASA (advanced germ cell marker in cyan), and SYCP3 (meiosis marker in magenta) and imaged with a 25X∼/0.95 water objective. On the top edge, there was non-specific signal for SYCP3 staining (edge effect). The optical section view presents the middle section of the ovary; asterisk indicates germ cell deprived area. In higher magnification, OCT4 negative; VASA and SYCP3 positive germ cells are bordered by dashed white lines. **C**. OCT4+ (yellow) and SYCP3+ (magenta) cells are shown as spots. Rectangle on the left shows the area where transverse optical section view is displayed on the right. SYCP3+ cells are in the middle of the ovary, near the mesonephric-gonadal junction (white arrows).

**Figure 2.**
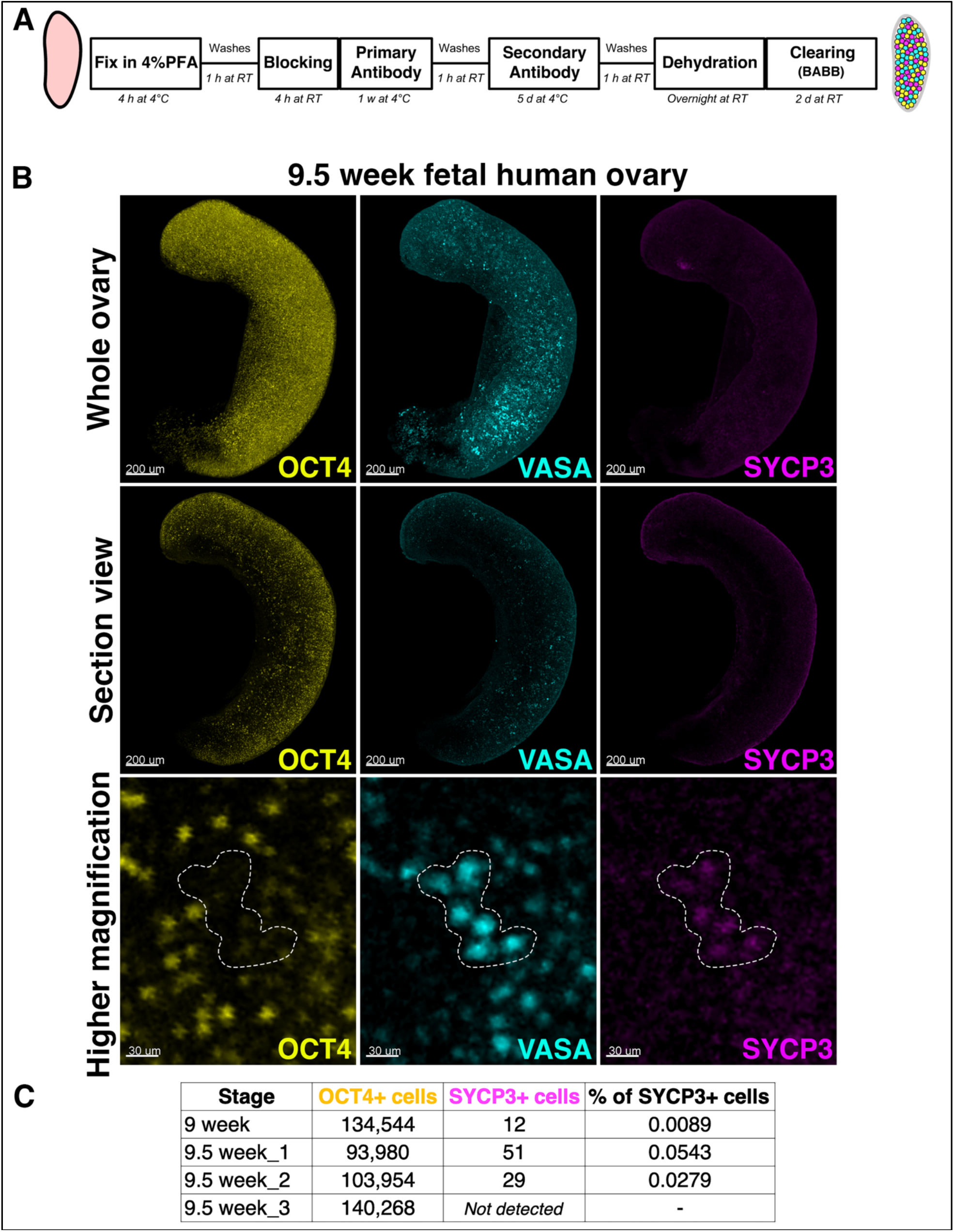
Quantification of pluripotent and meiotic germ cells in human fetal ovaries in 3D. **A**. Flowchart of whole-mount immunofluorescence labeling and organic solvent-based clearing (using BABB) of human fetal ovaries. **B**. Whole-mount image of 9.5-week-old ovary with 10X∼/0.40 dry objective. Higher magnification image of an OCT4 negative, VASA and SYCP3 double-positive germ cell cluster demarcated by dashed white lines. **C**. Quantification of OCT4+ (pluripotent) and SYCP3+ (meiotic) germ cells. In one ovary sample collected at 9.5 weeks, SYCP3+ cells were not detected.

By week 9.5, the number of meiotic SYCP3+ cells increased about three-fold compared to the 9-week specimen. Since our 3D analysis protocol relies on the nuclear signal for segmentation of objects, we were unable to quantify cytoplasmic VASA expression in fetal ovaries. The majority of SYCP3+ meiotic germ cells exhibited strong expression of VASA, whereas the pluripotency marker OCT4 was decreased in those germ cell clusters (nests) (**Figure 2B**). These results demonstrate that our 3D analysis technique enables the first quantification germ cells in intact human fetal ovaries, and identification of the first rare meiotic germ cells that are present in the center (interior) of the ovary at 9-weeks of development.

### After meiotic initiation, the number of meiotic germ cell is highly variable

At week 10 of development, SYCP3+ germ cell clusters remained in the center (interior) of the ovary and expanded in size and quantity (**Figure 3A**). The number of meiotic SYCP3-expressing cells increased in all 3 samples analyzed however, there was considerable variation between them (**Figure 3B**).

**Figure 3.**
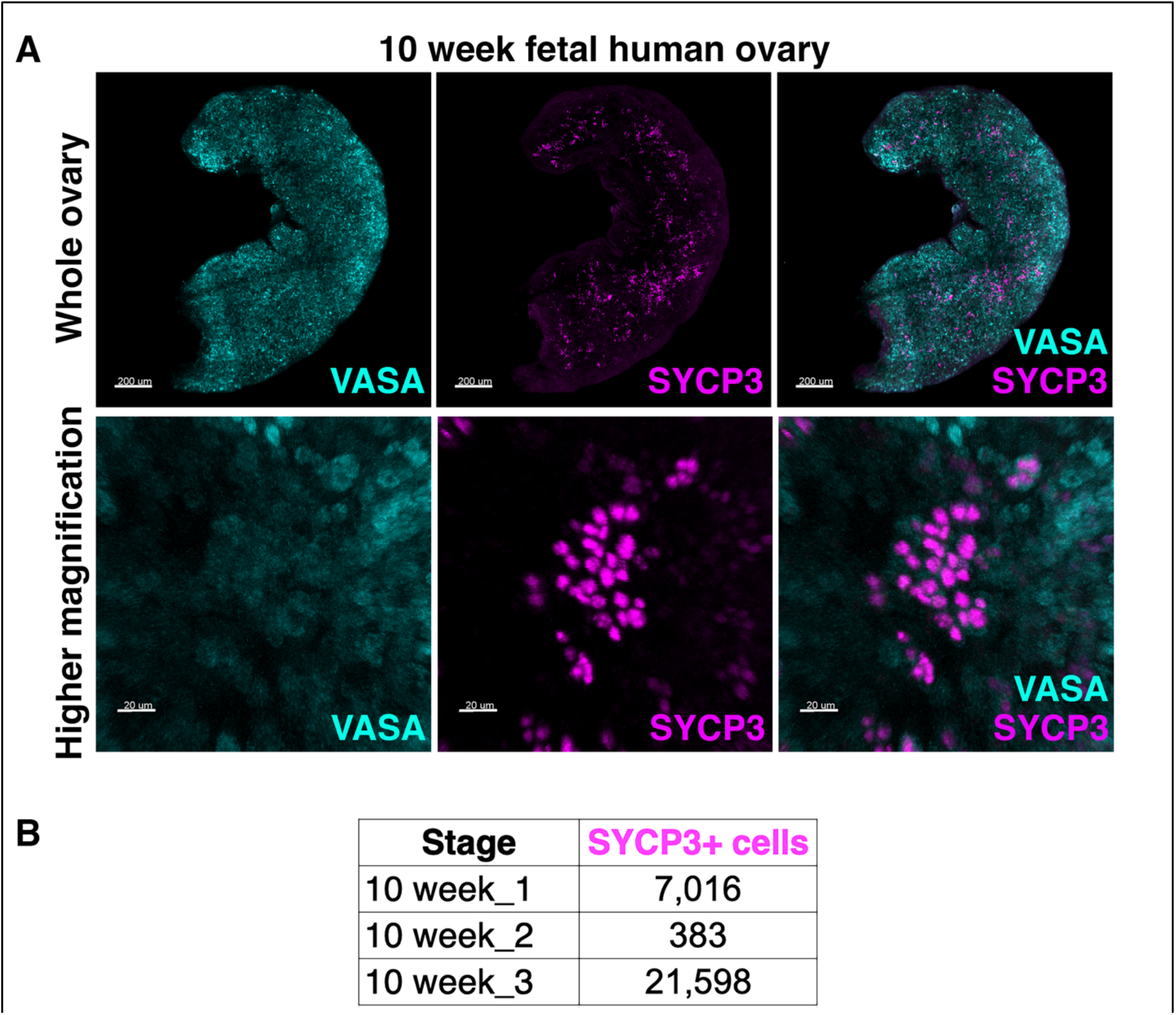
3D quantification of meiotic germ cells in 10-week-old fetal ovaries. **A**. An increased number of VASA and SYCP3-positive germ cells was observed at 10 weeks. Higher magnification images show an increase in the size of SYCP3+ germ cell clusters. It is important to note that most SYCP3+ cells are VASA+, but not all VASA+ germ cells are SYCP3+. **B**. Number of SYCP3+ cells in 10-weeks-old fetal ovaries.

A similar degree of individual variation in SYCP3+ cell numbers was seen at week 11 and week 12 (**Figure 4**). We observed two distinct meiotic germ cell populations; some SYCP3+ germ cells were OCT4 negative (**Figure 4A**), whereas others maintained OCT4 expression (**Figure 4B**). Due to the technical limitations, it remains unknown whether the two distinct meiotic germ cell populations are members of the same germ cell cyst (physically connected) or belong to the same clone (shared ancestry).

**Figure 4.**
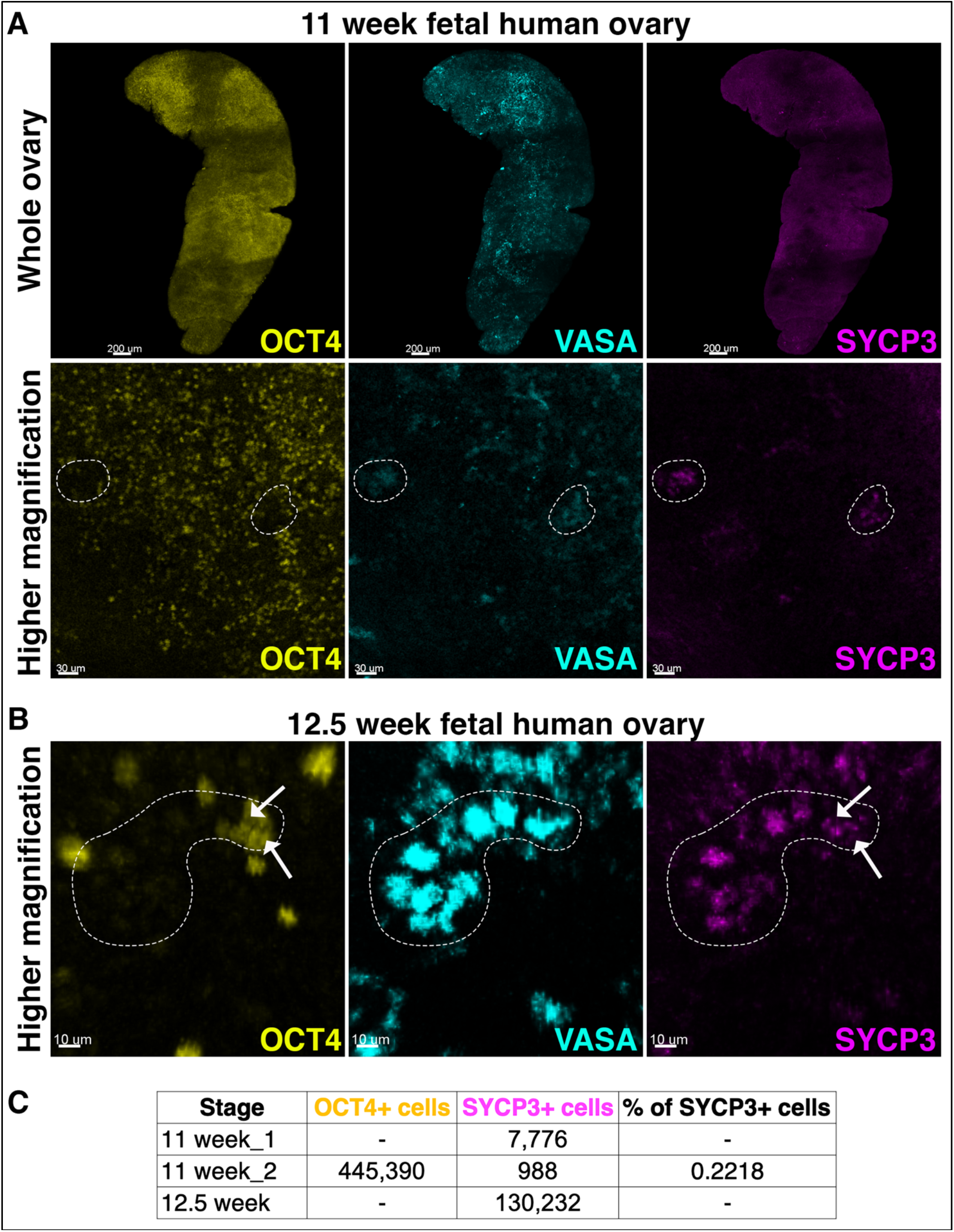
Asynchronous state transition in human fetal ovaries. **A**. Intact 11-week-old fetal ovary immunostained for OCT4 (yellow), VASA (cyan) and SYCP3 (magenta). Shown at higher magnification, the white dashed lines indicate a cluster of germ cells that are OCT4-negative, VASA and SYCP3 positive. **B**. VASA+ germ cells (in a cluster marked by dashed lines) are SYCP3+ and partially OCT4+, demonstrating less synchronous state transitions in some germ cells in the cluster. **C**. Quantification of OCT4+ (pluripotent) and SYCP3+ (meiotic) germ cells in 11-and 12.5-week-old fetal ovaries.

Despite the widespread expression of SYCP3 by 12.5 weeks of development, whole-mount imaging revealed the presence of SYCP3-negative germ cells at/near the surface of the ovary (**Figure 5A**). A comprehensive examination of first trimester human fetal ovaries from week 9 to 12.5 revealed a pattern of radial meiotic initiation, with clusters of SYCP3+ germ cells concentrated at the center (interior) of the ovary and progressively expanding outward (**Figure 5B**). Such a substantial degree of variation in the number of meiotic germ cells has been assumed but never directly observed.

**Figure 5.**
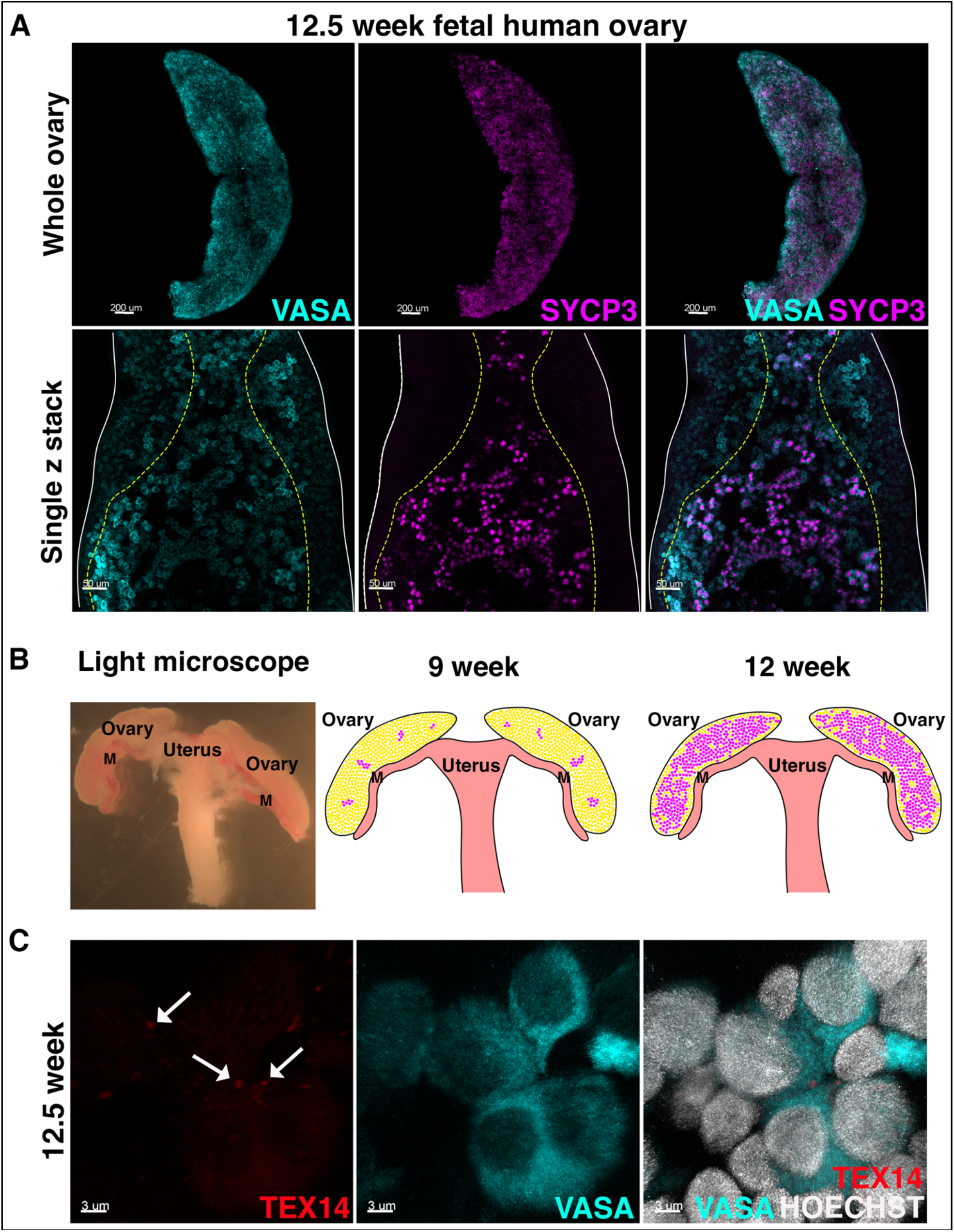
A pattern of radial meiotic initiation and intercellular bridges in human ovaries. **A**. SYCP3+ cells are absent from the periphery of the ovary at 12.5 weeks FA (yellow dashed lines in single z stack). **B**. A model of radial meiotic initiation in the human ovary. The yellow and magenta cells represent non-meiotic and meiotic cells respectively. M; mesonephros. **C**. Immunostaining in sections reveals TEX14+ foci at the borders between VASA+ cells at 12.5 week.

As meiosis in the mouse ovary initiates in germ cell nests (Lei & Spradling, 2013) through physical coordination across specialized intercellular bridges (Soygur et al., 2021), the observed clustering of SYCP3 expression suggested a similar mechanism in human fetal ovaries. Previous studies reported the presence of intercellular bridges between developing germ cells in human fetal ovaries by ultrastructure (Gondos et al., 1971; Ruby, Dyer, Gasser, & Skalko, 1970; Stegner & Wartenberg, 1963). Although many components of germ cell intercellular bridges have been identified, only Testis expressed 14 (TEX14) was found to be required in their formation by genetic knockout studies in mice (Greenbaum, Ma, & Matzuk, 2007; Greenbaum et al., 2006). *TEX14* gene expression was detected in both female and male fetal germ cells (Li et al., 2017), and TEX14 protein localization was documented in intercellular bridges of human testis (Greenbaum et al., 2007), however evidence is lacking in human fetal ovaries. Immunofluorescent staining of human fetal ovary sections revealed the presence of TEX14 protein in foci between VASA+ germ cells at 12.5 weeks FA (**Figure 5C**). This result raises the possibility that germ cell communication via intercellular bridges may underlie the initiation of meiosis in clusters in the human fetal ovary as in mice, although the function of intercellular bridges is difficult to test in humans due to challenges in recapitulating meiotic initiation *in vivo*.

### Whole-mount analysis of fetal ovaries from the second trimester

To further investigate spatiotemporal dynamics of meiotic and non-meiotic germ cells, we extended our 3D whole-mount approach to ovaries from the second trimester. As we observed at week 18.5 of fetal development, meiotic germ cells identified by the expression of SYCP3 and ɣH2AX were abundant but absent from the surface of the ovary (**Figure 6A**). We furthermore observed substantial numbers of OCT4+ SYCP3-germ cells in the periphery of the ovary **(Figure 6B)**. This observation is consistent with previous 2D histologic studies that reported OCT4+ pluripotent germ cells localized exclusively in the cortex (or surface/periphery) of second trimester fetal ovaries (Anderson et al., 2007; Stoop et al., 2005). However, in contrast to this prior study, we also observed OCT4+SYCP3-cells intermingled with SYCP3+ germ cells throughout the center of the ovary (**Figure 6Biii-iv**).

**Figure 6.**
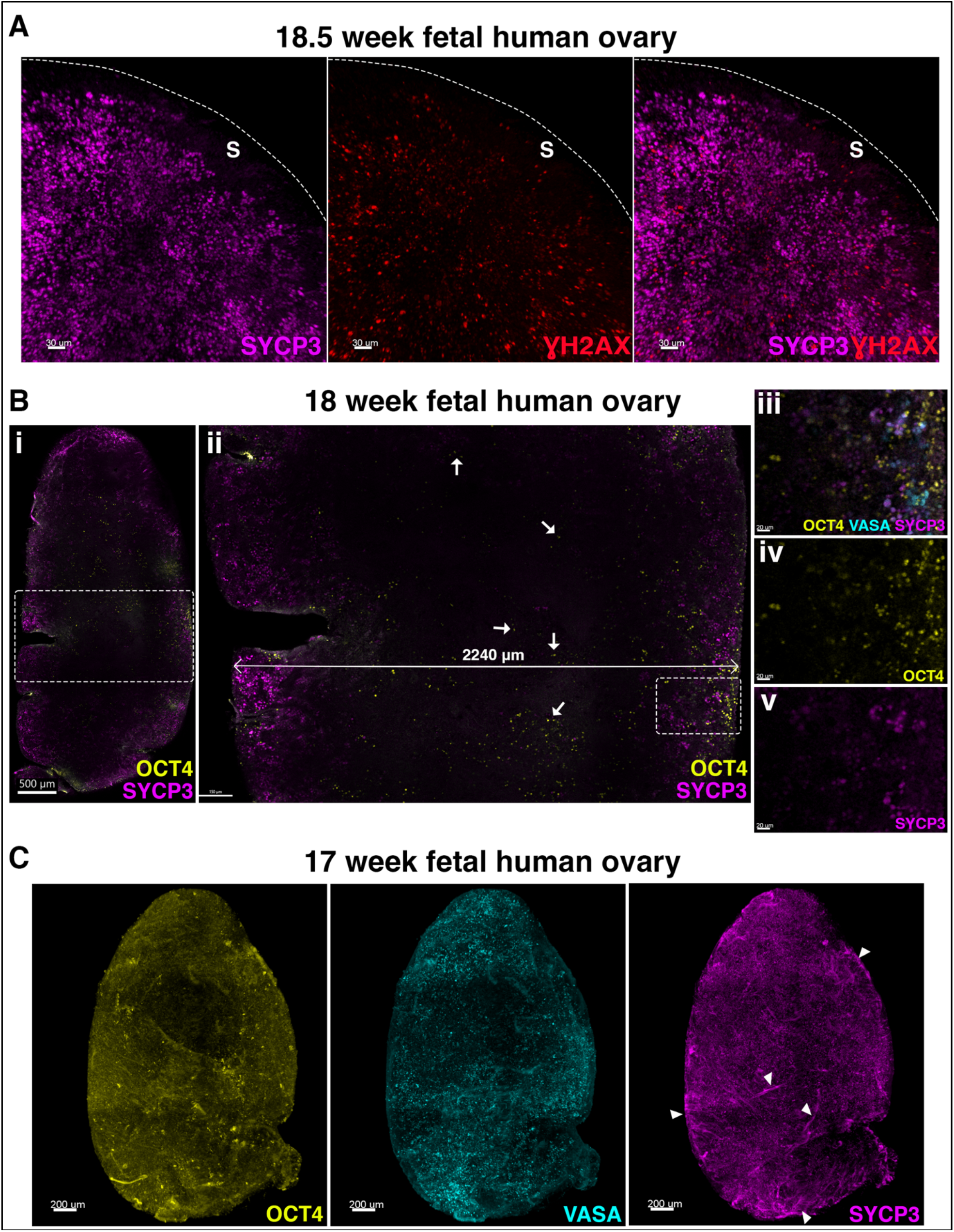
3D whole-mount analysis of second trimester human ovaries. **A**. Fetal ovary surface (S) lacking SYCP3 and ɣH2AX positive meiotic germ cells at 18.5 weeks of development. **B**. Optical section views of the fetal ovary at 18 weeks FA show OCT4+ cells throughout the ovary. Dashed rectangle in **Bi** represents the zoomed-in area in **Bii**. An ovary section towards the middle of the ovary measures 2240 μm in thickness and OCT4+ cells (in yellow) are scattered and visible throughout the entire section. A rectangle in **Bii** shows the zoomed in view of **Biii, Biv** and **Bv. C**. Whole-mount image of 17-week-old ovary, OCT4 in yellow, VASA in cyan, and SYCP3 in magenta. Arrowheads show non-specific binding of antibodies to connective tissue-like fibers.

In older ovaries (17-week and later), the accuracy of object detection was limited by the large size and complexity of the organ. Connective tissue-like fibers became visible (**Figure 6C, white arrows**) and non-specific antibody binding was observed. For these reasons, quantification of germ cells during the second trimester was less accurate than for ovaries from the first trimester. Despite suboptimal conditions, we detected around 75,000 OCT4+ cells and approximately 300,000 SYCP3+ meiotic germ cells at 18 weeks. This overall increase in SYCP3+ cells compared to first trimester ovaries (which peaked at ∼130,000 SYCP3+ cells at 12.5 weeks and ∼445,000 OCT4+ at 11 weeks) demonstrates progressive commitment of germ cells to meiosis during development.

## Discussion

Building 3D maps of human development requires the collective efforts of many individual labs as well as network initiatives. This study represents the first effort toward 3D modelling of the human fetal ovary at single cell level. Here, we describe the spatiotemporal characteristics of the mitotic-meiotic transition, providing a quantitative framework to analyze one of the major events in the human fetal ovary. Quantitative analysis revealed that 9 - to 9.5-week old ovaries harbor ∼118,000 OCT4+ germ cells which is almost twofold less than previously reported estimates from histologic sections (Bendsen et al., 2006). Using a synaptonemal complex protein3 (SYCP3) as a definitive marker of meiosis, we demonstrated the very first meiotic germ cells in the fetal ovaries at 9-weeks of development, earlier than previous histological (Le Bouffant et al., 2010), cytological and ultrastructural studies (Gondos et al., 1986). Given that the human fetal ovary grows from ∼one mm to four mms (in width) (Shen et al., 2018), a major limitation in studies using histological sections is the technical inability to quantify the absolute number of germ cells. The 3D quantitative approach we developed permits analysis of hundreds of thousands of germ cells in intact ovaries, detection of the first vanishingly small proportion of meiotic entrants, and visualization of anatomical relationships within the entire organ during this extended period of female germ cell sex differentiation.

Asynchrony of meiotic initiation has been shown in both mouse (Bullejos & Koopman, 2004; Menke, Koubova, & Page, 2003) and human (Baker, 1963; Kurilo, 1981; Skrzypczak et al., 1981) ovaries. A wave of meiotic entry was described in mouse ovaries that travels first from center to periphery (Soygur et al., 2021) and then from anterior to posterior (Bullejos & Koopman, 2004; Menke et al., 2003). The timing of this transition in mice has been attributed to several factors, including Retinoic Acid (RA) (Bowles et al., 2006; Koubova et al., 2006), epigenetic reprogramming via Tet1 and polycomb (Vincent et al., 2013; Yokobayashi et al., 2013) and to cytoplasmic sharing via intercellular bridges (Soygur et al., 2021). In human fetal ovaries, studies suggest that RA is partially responsible for regulation of the meiosis (Childs, Cowan, Kinnell, Anderson, & Saunders, 2011; Le Bouffant et al., 2010) however, the anterior-posterior (or cranial-caudal) wave is yet to be thoroughly studied. Using histological examinations to count germ cells in human fetal ovaries, Bendsen et al. found that early stages of meiotic germ cells could only be detected in the cranial part of the ovary near the rete ovarii, which connects the ovary to the mesonephros (Bendsen et al., 2006). Due to difficulties in maintaining anatomical orientation during the tissue obtaining process in our study, the cranial and caudal aspects of the ovary could not be definitively identified in our analysis. However, we observed that one extreme of the ovary had more SYCP3+ meiotic germ cells than the other at 9-10 weeks of age. Those first meiotic entrants tended to localize in the center (interior) of the ovary, and as meiotic frequency increased, this pattern expanded towards the periphery of ovary, resulting a radial wave of meiosis. Despite previous reports of developmentally advanced VASA-positive germ cells in the center of human ovaries, this germ cell marker is not related to meiotic entry. Using SYCP3 protein as a definitive marker of meiosis, we found that the radial initiation of meiosis occurs in human fetal ovaries in a way that is similar to mouse embryonic ovaries (Soygur et al., 2021), raising the possibility that human and mouse ovaries are more similar than previously thought. Surprisingly, we also observed that cells bearing the earlier PGC marker OCT4 were not restricted to the cortex but abundant throughout the center of the ovary even up until 18 weeks; this further underscores the asynchrony of human germ cell development.

Intercellular bridges (ICBs) are a unique feature of germ cell development that are evolutionarily conserved across many organisms from Drosophila to mice to humans. It has been well documented that ICBs transfer specific oocyte proteins, mRNAs, and organelles from supporting cells to the dominant oocyte in Drosophila [reviewed in (Greenbaum et al., 2011)]. In human fetal ovaries, ICBs were first characterized via electron microscopy (Ruby et al., 1970). The presence of ICBs within groups of degenerating germ cell suggested their potential role in this degeneration and ensuing engulfment by neighboring Granulosa cells (Gondos, 1973). Conversely, mice lacking ICBs suggested the opposite; ovaries deficient in the ICB component *Tex14* harbor fewer germ cells compared to wildtype ovaries at postnatal day 2 (PN2) (Greenbaum, Iwamori, Agno, & Matzuk, 2009). More recently, cytoplasmic sharing via ICBs was shown to be essential for appropriate timing of meiotic entry across the ovary and coordination of mitotic meiotic transition in mice (Soygur et al., 2021). Although we could not examine co-localization of SYCP3 and TEX14 in this study due to incompatibility of antibodies, the presence of TEX14 protein in developing germ cells supports the hypothesis that ICBs perform a similar coordinating role in human ovaries.

Given the emerging paradigm that human germ cell development proceeds from a pluripotent PGC stage to a VASA+ advanced germ cell stage and then to meiosis (Gkountela et al., 2013), an intriguing observation was the co-expression of OCT4 and SYCP3 in some germ cells, which may suggest a gradual and less synchronous change in cell state. A similar observation was presented by Anderson et al. (2007), who identified an intermediate population that expressed low levels of both OCT4 and VASA proteins in second trimester human ovaries. In that study, double-positive cells were located in the transition zone between periphery and center of the ovary (Anderson et al., 2007). In this context, it is surprising that we observed a substantial number of OCT4+ cells in the center of the ovary in even the second trimester of pregnancy. This result demonstrates the value of 3D imaging in identifying rare cell populations and spatiotemporal characteristics of germ cell development.

In conclusion, our work provides a roadmap for visualizing human fetal ovaries in 3D at the single cell level. Understanding 3D dynamics of early oocyte development in the ovarian microenvironment will help to uncover mechanisms governing mitotic-meiotic transition and eventually guide us to improve culture systems for *in vitro* oocyte derivation.

